# Morphology of Mitochondria in Spatially Restricted Axons Revealed by Cryo-Electron Tomography

**DOI:** 10.1101/273052

**Authors:** Tara D. Fischer, Pramod K. Dash, Jun Liu, M. Neal Waxham

## Abstract

Neurons project axons to local and distal sites and can display heterogeneous morphologies with limited physical dimensions that may influence the structure of large organelles such as mitochondria. Using cryo-electron tomography (cryo-ET), we characterized native environments within axons and presynaptic varicosities to examine whether spatial restrictions within these compartments influence the morphology of mitochondria. Segmented tomographic reconstructions revealed distinctive morphological characteristics of mitochondria residing at the narrowed boundary between presynaptic varicosities and axons with limited physical dimensions (~80 nm), compared to mitochondria in non-spatially restricted environments. Furthermore, segmentation of the tomograms revealed discrete organizations between the inner and outer membranes, suggesting possible independent remodeling of each membrane in mitochondria at spatially restricted axonal/varicosity boundaries. Thus, cryo-ET of mitochondria within axonal subcompartments reveals that spatial restrictions do not obstruct mitochondria from residing within them but limited available space can influence their gross morphology and the organization of the inner and outer membranes. These findings offer new perspectives on the influence of physical and spatial characteristics of cellular environments on mitochondrial morphology and highlights the potential for remarkable structural plasticity of mitochondria to adapt to spatial restrictions within axons.

## Introduction

Neurons are architecturally complex cells that can extend axonal projections with elaborate arborization for several hundreds of millimeters (and in some cases, meters) to form synaptic connections with local and distal targets [1–3]. Depending on the target, presynaptic compartments can either be found at the end of axons (terminal boutons) or tracking along axons as intermediate swellings (*en passant* boutons or varicosities), such as in the unmyelinated CA3->CA1 axons of the hippocampus [3–7]. The bead-like presynaptic varicosities are morphologically heterogeneous, displaying diameters that can range from 1-2 μm connected by thin axon segments that can have diameters less than 100 nm [4, 5, 8, 9].

Tracking through the axons is a well-developed system of microtubules that mediate motor-driven anterograde and retrograde transport of signaling cargoes, protein complexes and organelles critical for function and homeostasis at distant synapses [10, 11]. Transport of these intracellular components creates a spatially and temporally dynamic environment within the axon that can contain a variety of organelles and cargoes of different shapes, sizes, and number [12, 13]. As axon segments interconnecting varicosities can be remarkably thin and occupied by various structures, whether physical adaptations to available space are required for the motility of large organelles, such as mitochondria, with diameters ranging between 100-500 nm, poses an interesting question [14, 15]. Mitochondria within axons and synaptic compartments are particularly critical for development, function, and plasticity. During transport, mitochondria are known to make frequent stops, or saltatory movements, at presynaptic compartments to provide local ATP synthesis and calcium regulation required for proper neurotransmission [16–20]. Given the morphological complexity of axons, restricted physical dimensions and available space could potentially influence the subcellular localization, distribution, and transport of mitochondria required to meet local energy needs and for supporting synaptic transmission. Although mitochondria are morphologically dynamic organelles that can exist in a variety of shapes and sizes, how mitochondria adapt to the physical constraints presented in axons has not been previously examined.

Advances in microscopy and imaging techniques have played a pivotal role in revealing the three-dimensional architecture of neurons and their intracellular environments at resolutions reaching the nanometer scale [21, 22]. In the current study, we employed cryo-electron tomography (cryo-ET) to visualize three-dimensional spatial relationships and organellar structure within cultured hippocampal axons and varicosities. The unique native state preservation afforded by cryopreservation and the resolution of cryo-ET revealed that axon morphology and physically restrictive intracellular dimensions present a previously unrecognized influence on the morphology and ultrastructure of mitochondria residing at the boundary between large varicosities and small axonal subcompartments.

## Results

### Preparation of primary hippocampal neurons for cryo-ET

Primary neuronal cultures from E18 rat hippocampi were grown on holey carbon grids prior to cryo-preservation. At 10 days post plating, primary hippocampal neurons have extended dendritic and axonal processes and have established presynaptic varicosities and initial excitatory synaptic connections [23]. **Figure 1A** and **B** show representative bright field images of neurons cultured on a Quantifoil grid where widespread elaboration of processes is evident. **Figure 1C** and **D** demonstrate images of companion grids that were fixed and immunolabeled with antibodies to calcium/calmodulin dependent protein kinase II alpha (CaMKIIα) and synapsin 1 to visualize the excitatory neuron population and presynaptic varicosities, respectively. CaMKIIα antibodies largely label soma and processes, while the synapsin 1 antibody shows distinct puncta representing enrichment of synaptic vesicles at presynaptic varicosities.

**Figure 1.**
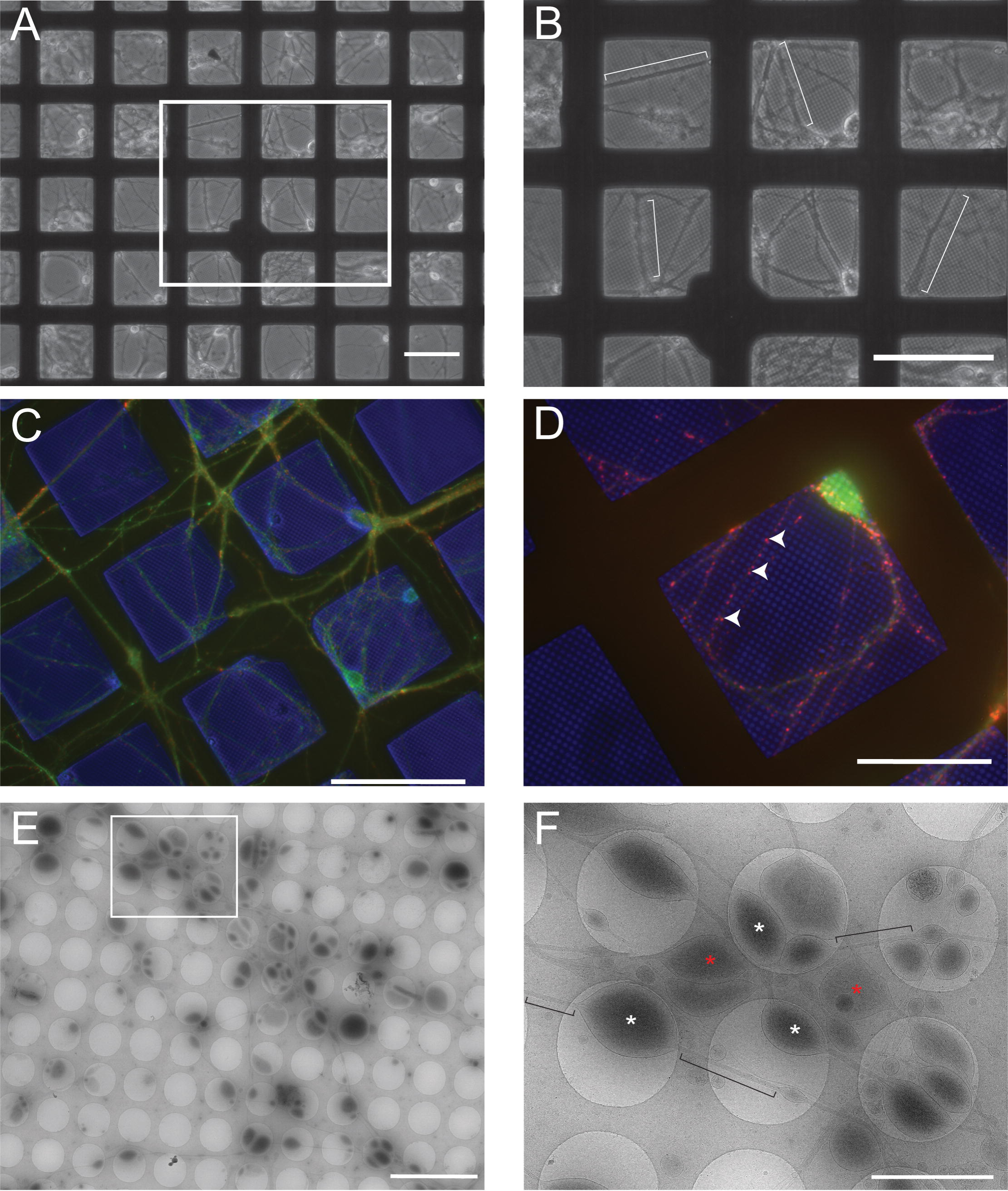
Growth and characterization of primary hippocampal neurons on EM grids. Hippocampal neurons were isolated from E18 rats and plated on poly-D-lysine coated Quantifoil 2/1 gold grids. A) Low magnification bright field image showing the typical distribution of neuronal soma and processes after 10 d in culture. B) Higher magnification image of the area from the white box in panel (A) with neuronal processes highlighted by white brackets. (C) Wide-field fluorescence image showing immunolabeling of the neuron specific protein CaMKIIα in green and the presynaptic vesicle associated protein synapsin 1 in red. The blue color is from a bright field overlay of the same area that also highlights the bars of the EM grid. (D) Higher magnification image in a different area of the same immunolabeled grid highlighting the punctate staining of synapsin I (red; arrowheads) along processes typical of *en passant* varicosities in hippocampal axons. Blue is again from where the grid bars and holes in the Quantifoil grid are apparent. Scale bars in panels A-C = 100 μm and in panel D = 50 μm. (E) Low magnification montage of one area in a cryopreserved grid of hippocampal neurons 10 d post-plating showing the typical distribution of axons and synaptic varicosities. Scale bar = 10 μm. F) Higher magnification representation from (E; white box) showing the axon and varicosity distribution overlying carbon (slightly darker areas) and grid holes. Examples of varicosities lying within grid holes are marked with white asterisks while examples lying on the carbon are marked with red asterisks. Axon segments interconnecting the varicosities are highlighted with black brackets. Scale bar = 2 μm.

For cryo-ET, fiducial gold markers were applied to prepared grids to aid in image alignment during data acquisition and image processing, and then cryopreserved by plunge freezing in liquid ethane. Cryopreservation conserves the near native state of the preparation permitting an assessment of spatial relationships between organelles present in different neuronal compartments free of fixation- or stain-induced artifacts. Low magnification images were first collected (**Fig. 1E**) and montaged to provide maps for targeting areas of interest for data collection. A higher magnification image reveals the distribution of presynaptic varicosities and axonal processes (**Fig. 1F**). Varicosities and axons can be seen residing on both the carbon and overlying the grid holes. To provide maximum contrast, tomographic data collection was targeted to cryopreserved structures within the grid holes. In these preparations, the increased thickness of the soma and proximal dendrites prevented sufficient electron beam penetration for imaging of these structures. In contrast, the sample thickness surrounding the axonal processes and varicosities was ideal, permitting a detailed assessment of the spatial relationships of cytoskeletal structures and organelles within these subcompartments.

### Organelle populations of presynaptic varicosities in primary hippocampal neurons are heterogeneous

Areas were randomly chosen, and tilt series were collected from varicosities and axon segments overlying grid holes to visualize cytoskeletal and organellar structures. **Figure 2A** shows a slice through a representative tomographic reconstruction with various resident organelles and structures visible, including two mitochondria, a multi-vesicular body (MVB), microtubules, endoplasmic reticulum (ER) and a collection of vesicles. Supplementary **Figure S1** shows 2D images of identified organelles and other structures observed within the population of varicosities analyzed. To determine the three-dimensional (3D) relationship between the different structures, segmentation was accomplished of the tomographic reconstruction of both the presynaptic varicosity and the adjoining axon (**Fig. 2B; Movie S1**). The reconstruction demonstrates microtubules (light blue) forming a continuous set of tracks traveling from one end of the varicosity to the other. Microtubules are well organized and relatively straight in axons, however, they exhibit greater curvature in the varicosity, while again gathering together and straightening when passing through the adjoining axon segment. The ER (yellow) also forms a reticulated and continuous structure spanning the entire length of the varicosity, consistent with other reports on the ubiquitous presence of ER in axons and synaptic terminals [21]. Mitochondria, MVB, and other sac-like compartments, exhibit more random distributions within the varicosity, while vesicles appeared to be somewhat clustered. To determine if the presence of each of these identifiable organelles was consistent across varicosities, we analyzed their distribution and characteristics in an additional 77 tomographic reconstructions. ER and microtubules were present in 100% of presynaptic varicosities, vesicles were present in 97%, mitochondria in 82%, sac-like compartments in 38%, MVBs in 21%, lamellar bodies in 9%, and autophagosomes in 4% (Table 1).

**Table 1.**
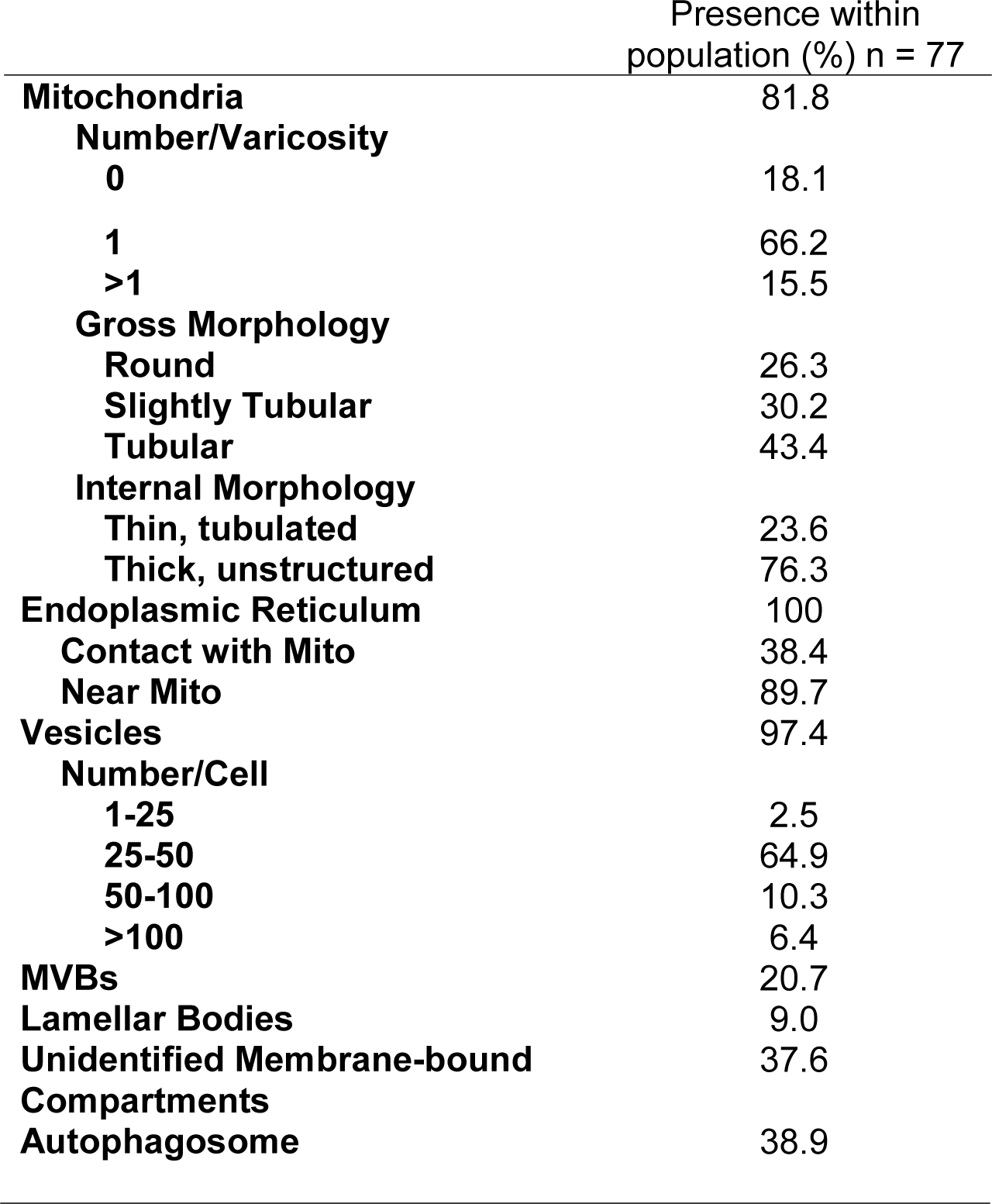
Organelle population representations

**Figure 2.**
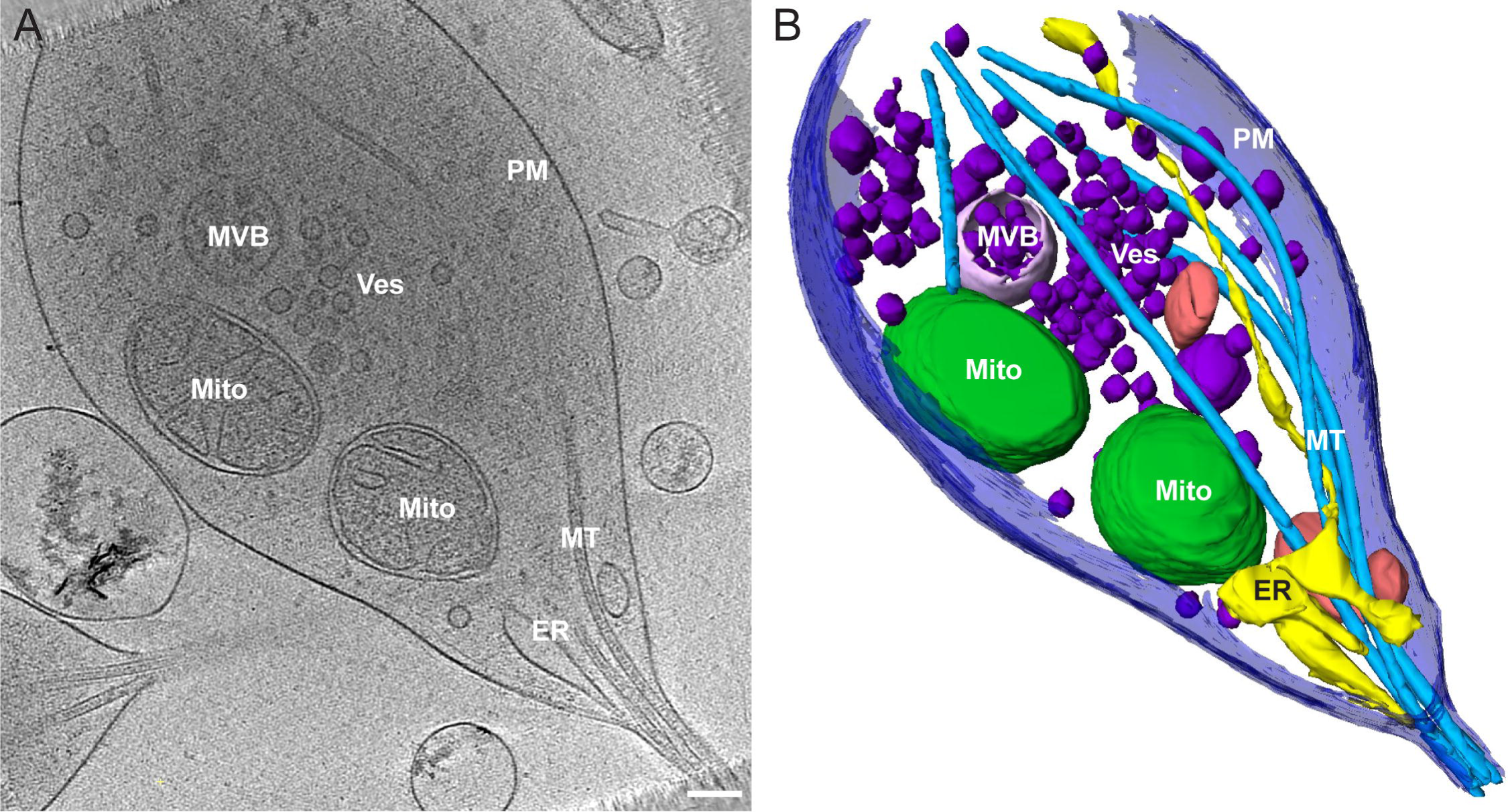
Tomographic reconstruction of a typical pre-synaptic varicosity and adjoining axon segment. A) 2D slice from the tomographic reconstruction showing the distribution of organelles in the varicosity PM: plasma membrane, MT: microtubules, Mito: mitochondrion, ER: endoplasmic reticulum, Ves: vesicle, MVB: multi-vesicular body. Scale bar = 200 nm. B) Segmented representation of the entire 3D tomogram volume shown in (A) revealing the relative size and spatial distribution of the organelle environment in the varicosity and axon segment. Plasma membrane (dark blue), microtubules (light blue), mitochondrial outer membrane (dark green), endoplasmic reticulum (yellow), vesicles (dark purple), multi-vesicular body (light purple), unidentified membrane-bound compartment (pink). Scale bar = 200 nm.

Variations in distinct sub-features of the observed organelle populations also emerged in 3D at high resolution. Specifically, the ubiquitous ER was observed in close apposition to almost every other organelle within the varicosities, consistent with ER-membrane contact sites described by others [21]. The vesicular population was heterogeneous in number, ranging from 1 to >100 vesicles per varicosity. 75% of varicosities contained <50 vesicles, 10% had 50-100, and 6% contained more than 100 (Table 1). Vesicles were observed in clusters with electron-dense filamentous connections (data not shown), consistent with previous reports of proteinaceous (synapsin, Bassoon, and/or ERC2) tethering of synaptic vesicles [12, 13]. Mitochondrial cristae structure within the 3D tomographic datasets was variable, but could be broadly segregated into two distinct populations based on ultrastructural features (**Fig. S2**). A population of mitochondria displayed the canonical thin, tubular cristae morphology (**Fig. S2A**), while a second population displayed thick, unstructured cristae (**Fig. S2B**).

In contrast to the heterogeneous appearance and distribution of organelles in presynaptic varicosities, the adjoining axonal segments were more consistent in composition. The most obvious components were microtubules that were seen as continuous elements, gathered at the sites of axonal narrowing at both ends of the varicosity. Microtubules did not display variability in diameter (~20 nm), however they did vary in number among different processes and occupied a significant portion of the available volume within the axon. As noted, microtubules are the essential tracks required for motor driven organelle transport and are critical for maintaining synaptic homeostasis and neuronal signaling. The corresponding microtubule occupation of axonal space leaves the qualitative impression that the available volume in axons to accommodate large organelles, such as mitochondria, might be an under-appreciated constraint affecting transport. The magnitude of this spatial constraint can be visually appreciated in **Movie S2**.

### Mitochondria display distinct morphological features at spatially restricted axon/varicosity boundaries

Previous EM studies have reported an average axonal diameter between 0.08 and 0.4 μm for unmyelinated cortical axons and an average varicosity diameter of ~1-2 μm [1, 5, 7, 8]. Varicosities and axons in our cryopreserved hippocampal preparations exhibited slightly smaller Feret diameters, with an average of 649 and 81 nm for varicosities and axons, respectively (Table 2). Additionally, the average Feret diameter of mitochondria observed in hippocampal varicosities was ~250 nm. These dimensions (summarized in **Table 2**) further reinforce the idea that significant spatial constraints exist that may influence mitochondrial structure in axons.

**Table 2.**
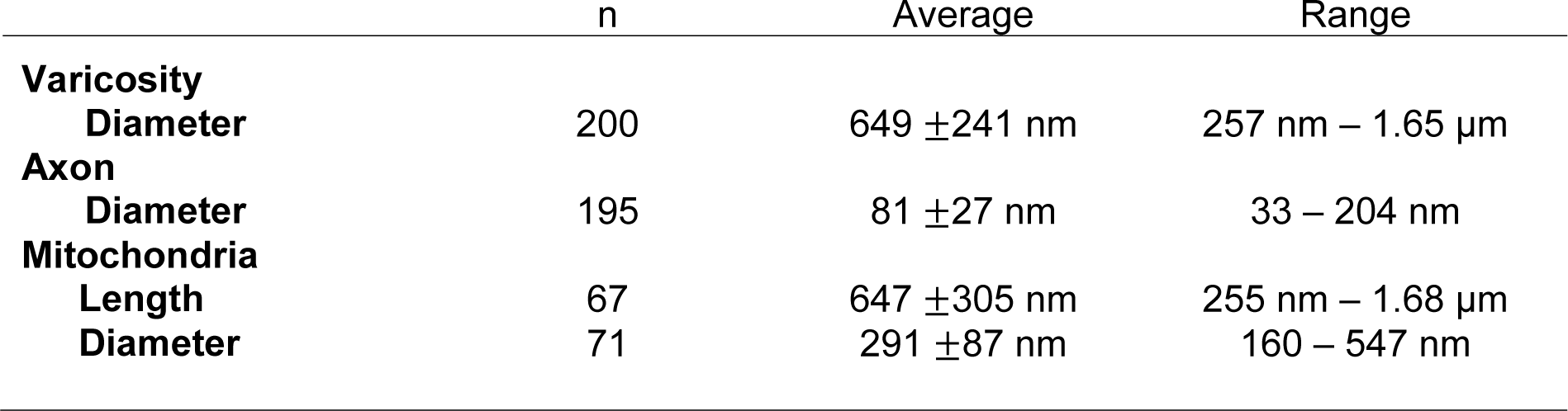
Two-dimensional morphological measures

For more in-depth investigation of this issue, high magnification tomographic data were collected targeting mitochondria residing at the boundary where the varicosity narrows into the small axonal segment. While such precise mitochondrial positioning was rare in the cryo-preserved neuronal population, the static events that were captured revealed distinct mitochondrial morphologies at the boundaries between varicosities and axons. **Movie S3** and **S4** demonstrate a mitochondrion partially residing in both a neuronal varicosity and the adjoining axon segment among the other organelles occupying space in these compartments. The portion of the mitochondrion residing in the varicosity is 305 nm in Feret diameter, while the portion of the mitochondrion residing in the axonal segment is narrowed to only 70 nm in diameter at its tip. Thus, the mitochondrion displays a major morphological change with an approximately 77% reduction in diameter within the axon. **Figure 3A** shows a snapshot of the segmented mitochondrion as well as additional examples of mitochondria captured at the varicosity/axonal boundary (**Fig. 3B and C**). **Figure 3C** shows a reconstruction that revealed two mitochondria residing in the varicosity and adjoining axon segment, with a portion of a third residing mainly in the axon. **Movies S3 – S10** demonstrate several tomographic reconstructions of mitochondria displaying similar morphologies at the varicosity/axon boundary. **Figure 4** and **Movie S9** show a particularly revealing example, in which a mitochondrion was captured bridging a short (100 nm in length) axon segment between two varicosities. The short narrow space produced a barbell shaped mitochondrion with a diameter between 200-300 nm in both varicosities while the portion spanning the axon segment was only 19 nm in diameter. To illustrate the ultrastructural features of the mitochondrion spanning the two varicosities, the inner mitochondrial membrane (IMM) and cristae were segmented in addition to the outer membrane (OMM; **Fig. 4C**). The resolution of this tomographic reconstruction was not sufficient to determine whether the IMM was continuous through the short axon segment, however **Movie S10** demonstrates an additional example of a mitochondrion spanning a short axon segment between two varicosities, in which the IMM appears continuous in the constricted section of the axon. Analysis of eight mitochondria exhibiting these drastic morphological features revealed an average of 84% (n = 8, SD = 6%) reduction in the Feret diameter of the mitochondrial area in the varicosity relative to the adjoining axon segment. These captured events highlight the potential adaptability of mitochondrial morphology to accommodate the available space within an axon. Note that in all of the segmented tomographic reconstructions in **Figure 3 and 4**, the presence of microtubules and additional organelles, such as the ER, further restricts the available volume to mitochondria within axons.

**Figure 3.**
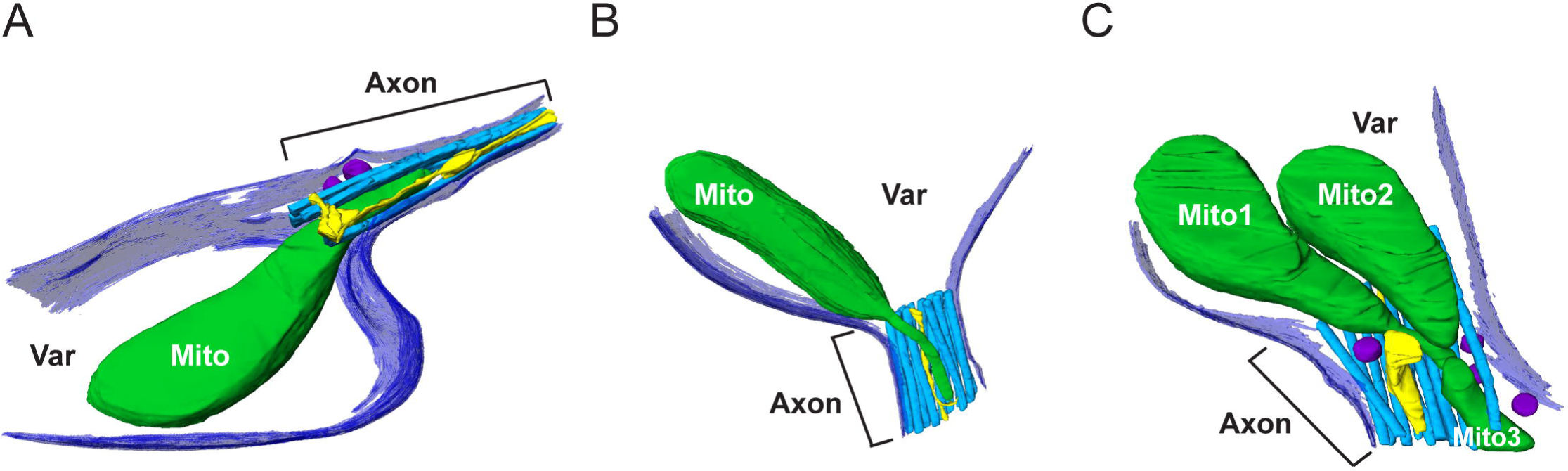
Mitochondria display atypical morphological features in physically restrictive axons. A-C) Three different 3D segmented reconstructions showing representative examples of mitochondria residing partially in the varicosity and adjoining axon segments, demonstrating different morphological states at the transition from the varicosity into the restricted space of the axon segment. Additional organelles occupying the axon space are also segmented. For ease of visualization, not all of the organelles and structures in the varicosity are shown. The plasma membrane (dark blue), microtubules (light blue), mitochondrial outer membrane (dark green), endoplasmic reticulum (yellow), vesicles (dark purple) are highlighted.

**Figure 4.**
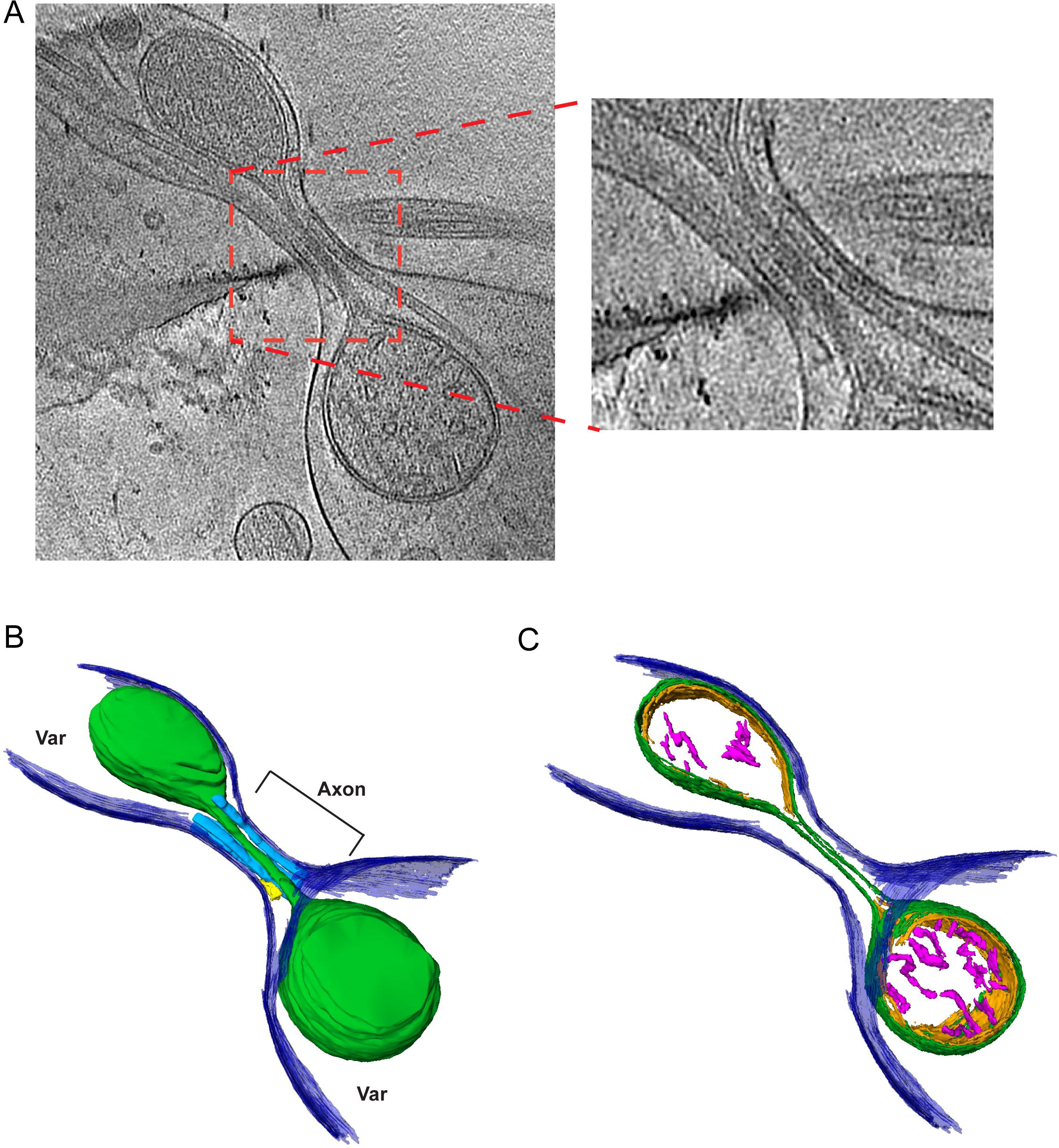
Structural features of a mitochondrion captured spanning two varicosities. A) 10 nm slice through a 3D tomographic reconstruction showing a mitochondrion spanning two closely spaced varicosities connected by a short (~100 nm) axon segment. An expanded region of the red box shown in (A) reveals the tubulated nature of the portion of the mitochondrion within the axon segment. Microtubules can be seen running parallel to the tubulated portion of the mitochondrion. B) shows a surface rendered version highlighting the plasma membrane (purple), microtubules (blue) small segment of ER (yellow) and mitochondria (green). C) is the same mitochondrion as in (B) displaying distinct segmentation of the outer membrane (green), inner membrane (orange), and cristae (pink).

### Mitochondrial membranes display unique organizations within spatially restrictive axons

Mitochondrial ultrastructure is thought to be dynamic with remodeling of the inner membrane and formation of specified subcompartments, termed cristae [24–26]. Given the atypical gross morphologies of mitochondria we observed between varicosities and the adjoining axonal segments, we questioned if the inner mitochondrial membrane also displays distinct structural features in axons with limited physical dimensions. To address this issue, the outer and inner membranes of mitochondria captured at the boundary between varicosities and adjoining axons were segmented, separating the outer membrane, and two regional components of the IMM, the inner boundary membrane (IBM) and the cristae. A conservative approach was taken during manual segmentation of cristae (i.e., only clearly discernible cristae membranes were included). Segmentation of each of these mitochondrial components revealed discrete morphologies between the inner and outer membranes. Most notably, two distinctions were observed between the outer membrane and the adjacent IBM at the narrowed mitochondrial tip in the adjoining axon segment. First, in some instances the IBM was observed to maintain apposition to the outer membrane at the narrowed mitochondrial tip entering the axon (**Fig. 5A**). Both the inner (orange) and outer (green) membranes can be seen narrowing as they enter the restricted axonal space. Interestingly, cristae (pink) are largely absent from this narrowed portion of the mitochondrion (~69 nm in diameter) residing in the axon. Second, the outer membrane was observed to separate from the inner membrane, leaving a space free of the inner membrane and matrix of the mitochondrion (**Fig. 5B**). Thus, it appears the IMM does not always remain in apposition with the OMM within the rather dramatic tubulation evident of the outer membrane residing within the restricted space of the axon. Two out of the eight representations of mitochondria displaying these atypical morphological features in our dataset also show the OMM separated from the IMM. **Movie S11** shows the segmented model of the three mitochondria in **Figure 3C and 5C**. The left mitochondrion can be seen to display the OMM separated from the IMM, while the right mitochondrion displays the OMM and IMM in juxtaposition, demonstrating the occurrence of both events in one varicosity/axon boundary area.

**Figure 5.**
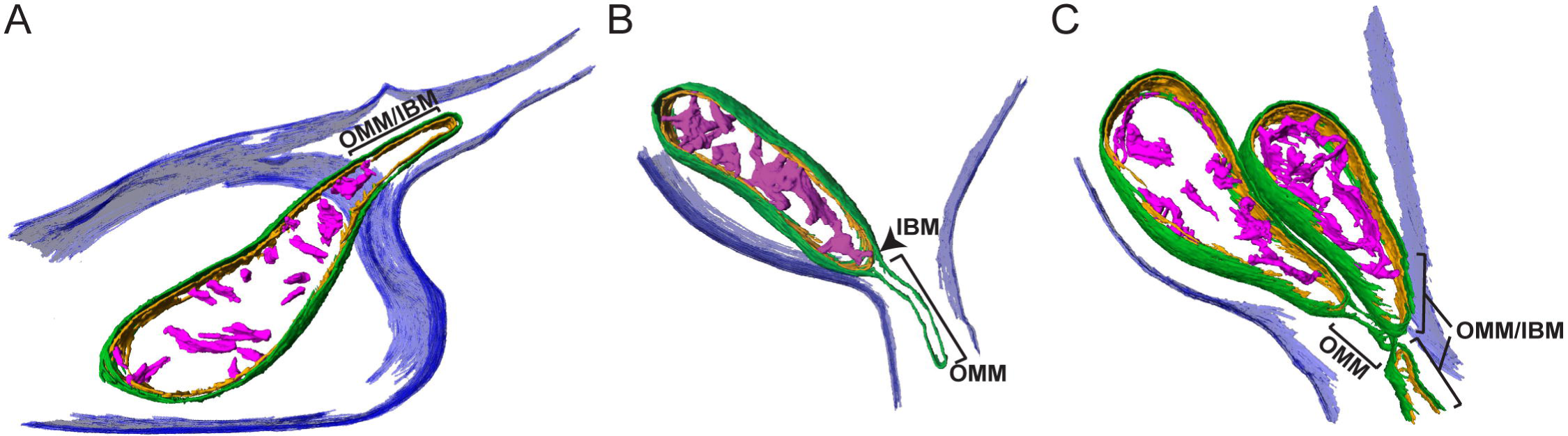
Mitochondrial membranes display distinct morphological features at the boundaries of varicosities and axons with limited physical dimensions. A-C) To highlight the membrane organization of the mitochondria shown in Figure 3A-C, the cristae and inner and outer membranes of the 3D reconstructions were segmented. A) Mitochondrial inner and outer membranes remain in close apposition within the narrowed portion of this mitochondrion resident in the axon. B) Mitochondrial outer membrane is separated from the inner membrane, creating a distinct “matrix-free” compartment in this portion of the mitochondria residing in the axon. C) Three mitochondria near the varicosity/axon junction show distinct internal membrane organization. The left mitochondrion shows a portion of outer membrane separated from inner membrane while the top right mitochondrion is narrowed near the axon junction, but the inner and outer membranes remain in apposition. A short tip of a third mitochondrion is partially captured at the edge of the tomogram that also shows inner and outer membranes together. Plasma membrane (dark blue), mitochondrial outer membrane (dark green), mitochondrial inner boundary membrane (orange), cristae (pink).

## Discussion

Although the transport of mitochondria within axons has been widely studied, the potential morphological adaptation of these large organelles to restricted physical dimensions and available space within axons has not been discussed [18, 27, 28]. The current study provides insight into spatial environments within presynaptic varicosities and thin axons of cryopreserved hippocampal neurons, unperturbed by fixation or stains via 3D cryo-ET. Distinct morphological characteristics of mitochondria were revealed at the boundaries between large varicosities and axon segments with limited physical dimensions (~ 80 nm). To our knowledge, this is the first study to describe such atypical mitochondrial morphologies apparently influenced by the limitations of physical space within thin axons. Additionally, the 3D reconstruction and segmentation of mitochondrial ultrastructure revealed distinct morphological features between the inner and outer mitochondrial membranes at spatially restricted axonal/varicosity boundaries, suggesting possible differential regulation of each membrane during these morphological adaptations.

Axon morphology can be widely variable depending on brain region. *En passant* boutons are common in axons of the hippocampus and cortex, giving rise to heterogeneous axon morphologies with presynaptic varicosities distributed along thin unmyelinated axons [3–7]. As axons and synaptic varicosities are dynamic, heterogeneous environments that can be occupied by organelles and molecules differing in size and number, it is important to consider whether available space within the axon may constrain motility or transport of large cargo, such as mitochondria. Using the advantages of cryo-ET, we were able to capture the static spatial environments in thin axons and presynaptic varicosities of cultured hippocampal neurons to examine organelle characteristics and distribution. Mitochondria, in particular, at an average of 250 nm in diameter, presented a clear spatial challenge to inhabit axons that are, on average, three times smaller (~80 nm). In the 3D segmentation of mitochondria residing at the boundary between the larger presynaptic varicosity and thin adjoining axon segments, we observed mitochondria displaying atypical morphological features. Mitochondria displayed a normal morphology within the varicosity and a narrowed tubulated portion, creating a “tip” that existed in the narrowed space within the axon. Interestingly, in some cases this narrowed portion of the mitochondrion was smaller than the inner boundary of the axonal plasma membrane, suggesting that additional material, not apparent in the tomograms, might further constrain the available space within the axon. The ability of mitochondria to display morphological diameters near 20 nm when challenged with limiting available space within axons is surprising and suggests mitochondrial morphology may be more adaptable in nature than previously considered. While it is important to emphasize that our methodology does not address temporal dynamics of mitochondria within axons, these observations highlight the potential for the adaptability of mitochondrial morphology and present interesting questions to the mechanisms that may be involved.

Movement of Intracellular cargo and transport within axons is mediated by microtubule-associated motor proteins that create the driving force necessary to pull organelles through the cytoplasm in axons and varicosities [11, 28]. Specifically, kinesin and dynein motor proteins exert a mechanical force on the mitochondrion to drive polarity-directed movement within axons through interactions with outer mitochondrial membrane and adaptor proteins, such as Miro and Milton [29]. We speculate that directional forces induced by motor proteins may drive the morphological features of the mitochondrial membrane, as observed in the current study. Kinesins, in particular, are known to induce membrane deformation or tubulation in *in vitro* reconstituted membranes [30, 31]. A recent study also described similar thin tubulation of mitochondria that is mediated by KIF5B, a member of the kinesin family [32]. Interestingly, the influence of mechanical forces on mitochondrial membrane dynamics was recently demonstrated by the recruitment of mitochondrial fission machinery and subsequent division at sites of induced physical constriction [33]. In fact, it is also possible that recruitment of such machinery would lead to the production of fission intermediates of the mitochondria of reduced size that would facilitate their movement through axons. If so, appropriate machinery would have to be present in adjacent varicosities for the reassembly of mitochondria. Although the distinct morphological features of mitochondria in axons observed in our static, cryopreserved tomographic reconstructions cannot address dynamics for transport, whether force-driven microtubule interactions play a role is an interesting possibility. Additionally, it is also possible that neuronal activity might influence the structure of varicosities, axons or mitochondria that would impact the magnitude of this problem. In this context, a recent report analyzing varicosities and axons in hippocampal slices, showed that high-frequency stimulation of axons, increased the size of varicosities and axons [34], although the peak effects on size were relatively modest (~5% increase in varicosity/axonal diameter). Thus, further investigation into the precise mechanisms involved in the regulation of space within axons and mitochondrial morphological adaptations within the available space is warranted.

Mitochondrial function is highly dependent on the unique architecture of the inner mitochondrial membrane (IMM), in which the respiratory complexes along with ATP synthase are concentrated in formations of distinct compartments, termed cristae [26, 35–37]. Classic and more recent studies examining inner membrane morphology and cristae formation have shown distinct ultrastructural features in states of high cellular energy demands, however whether inner membrane structure changes occur correspondingly with gross morphological changes (excluding those involved in mitochondrial fission or fusion) is not well defined [24–26, 38–41]. Therefore, we questioned whether the significant reduction in size of mitochondria within the physically restrictive axonal space also affected their internal structure. Segmentation of cristae and the inner boundary membrane (IBM), distinct components of the IMM, revealed intriguing differences in the relationship between internal structure and the outer mitochondrial membrane (OMM). In some cases, the IBM remained in close apposition with the OMM at the narrowed tip of the mitochondrion in the axon. Interestingly, this area was also void of cristae, whereas cristae structure remained unperturbed in the portion of the mitochondrion that resided in the varicosity, suggesting possible differential regulation of cristae compartments with adaptation to the limited space within axons. Conversely, a few mitochondria displayed a dissociation of the inner and outer mitochondrial membrane, in which the outer membrane was no longer in apposition with the IBM, but distinctly separated in the narrowed tip of the mitochondrion creating a “matrix-free” space. The IBM and cristae remained intact and unperturbed within the portion of the mitochondrion residing in the varicosity. The visualization of the 3D ultrastructure of mitochondria displaying these morphological features within spatially restricted axons suggests a potentially new instance of structural remodeling of the IMM. Mechanistically, this observation generates several interesting questions to the regulation and dynamic nature of mitochondrial ultrastructure, as well as the functional status of mitochondria in different axonal compartments. Although mechanisms for regulating inner membrane morphology remain incompletely defined, recent research has provided insight into some mechanisms involved in the regulation of cristae structure and IMM morphology [25, 37, 42, 43]. In addition to Opa1, the primary IMM fusion protein, ATP synthase and the mitochondrial contact site and cristae organizing system (MICOS) exhibit membrane bending functions and are proposed to determine curvature of the cristae membrane [37, 38]. MICOS components have also been implicated in regulating cristae morphology, as well as inner-outer membrane tethering, which could be a potential mechanism driving the differences in inner-outer membrane apposition or dissociation observed in the present study [44, 45]. Moreover, distinct events of IMM constriction independent from the OMM have also recently been observed that are driven by increased mitochondrial matrix calcium levels [46–48]. Thus, there is some evidence at the cellular level for differential morphological regulation of the inner and outer membranes. The 3D reconstructions of the OMM, IBM, and cristae in the cryo-preserved axon, and the observed dissociation between membrane morphologies in the current study lends support for the notion that the OMM and IMM can be remodeled independently. Functionally, the observed changes in cristae and IMM morphology also pose the question as to whether the lack of cristae or adaptation of the IMM influence the functional capacity of these mitochondria. Although the technical limitations of measuring mitochondrial function in relation to mitochondrial structure are challenging, these will be critical investigations in uncovering the dynamic and adaptive nature of the IMM and regulation of mitochondrial function in axons.

The importance of mitochondrial dynamics and synaptic localization to support neuronal function is well recognized (for reviews see [17, 18, 49]). Mitochondria must traffic through axons and populate distal synapses to mitigate local energy depletion and maintain calcium homeostasis required for vesicle release and recycling [16, 50, 51]. Additionally, mitochondria undergo active fission, fusion, mitophagy, and maintain contact with other organelles to communicate, functionally adapt, and maintain quality control within the local environment. Thus, disruptions in mitochondrial motility and dynamic behaviors can be detrimental to synaptic communication, plasticity, and survival, and have been implicated in several neurodegenerative processes (reviewed in [17, 29, 52]). The current study highlights the potential implications for spatial restrictions dictated by axon morphology to influence mitochondrial morphology, and potentially motility within the axon. These findings offer new perspectives on the physical and spatial influence of the cellular environment on mitochondrial morphology and highlights the remarkable structural plasticity of mitochondria to adapt to the limited available space within axons. Given the importance of membrane structure for mitochondrial function and the necessary transport of mitochondria for maintaining synaptic health and neurotransmission, these findings have far reaching implications for mitochondrial and neuronal biology. Future research will be essential to gain mechanistic insight to regulation of structural changes at mitochondrial membranes and the influence of morphological adaptations of mitochondria during axonal transport.

## Materials and Methods

### Neuronal Culture and Cryopreservation

Primary neuronal cultures were prepared from E18 rat hippocampi. All protocols involving vertebrate animals were approved by the Institutional Animal Care and Use Committee prior to initiating the studies. Briefly, pooled hippocampi were digested with papain for 20 min at 37°C and then triturated with a 10 ml pipet. Cells were counted and diluted in Opti-MEM containing 20 mM glucose to a density of 1.5 × 10^5^/ml. 200 mesh gold grids covered with Quantifoil 2/1 carbon were placed in 35 mm glass bottom Mat-Tek dishes and were treated overnight with 100 ug/ml poly-D-lysine. The dishes were then washed 3x with sterile water before plating cells at a density of 1.5 × 10^6^/dish. After letting the cells attach for 1 hr at 37°C/5%CO_2_, the media was exchanged for Neurobasal A supplemented with 2% B-27 (Life Technologies), GlutaMAX (Thermo Scientific, Waltham, CA), and penicillin-streptomycin (Sigma, St. Louis, MO) and incubated for 10 d at 37°C/5%CO_2_.

To cryopreserve intact neurons, the grids were lifted from the Mat-Tek dishes and 5 uL of Neurobasal media containing BSA coated 10 nm gold fiducials was applied. Fiducial gold facilitates tracking during image acquisition of tilt series and alignment of image frames during post-acquisition processing. After manual blotting, the grids were plunged into liquid ethane cooled with liquid N_2_. The entire process between removal of the grid from the culture dish and plunge freezing was on average ~30 s, but never more than 60 s. Cryo-preserved grids were stored in liquid N_2_ until use.

### Immunocytochemistry

Immunostaining of neurons grown on Quantifoil grids was accomplished by fixing the neurons at 10 d post-plating in freshly prepared 4% paraformaldehyde in 0.1 M phosphate buffer, pH 7.4, for 10 min at room temperature. The fixative was removed and reaction quenched with a 5 min incubation in 50 mM glycine in 0.1 M phosphate buffer, pH 7.4. Neurons were permeabilized with 0.5% TX-100 in 0.1 M phosphate buffer, pH 7.4, for 15 min and then non-specific sites blocked with Blocking buffer (2% normal goat serum, 1% bovine serum albumin, 0.1% TX-100 in 0.1 M phosphate buffer, pH 7.4) for 30 min. Primary antibodies were diluted 1:1000 in Blocking buffer and incubated for 1 hr at room temp. Primary antibodies included a monoclonal antibody to CaMKIIα (created in our lab; [53]) and a rabbit polyclonal antibody to synapsin 1 (Synaptic Systems Inc.). Grids were then washed 3x, 5 min each, with Wash buffer (0.2% normal goat serum, 0.1% bovine serum albumin, 0.01% TX-100 in 0.1 M phosphate buffer, pH 7.4). Grids were then incubated in 1:500 dilution of Alexa 488 labeled goat anti-mouse IgG and Alexa 568 labeled goat anti-rabbit IgG diluted in Blocking buffer for 30 min at room temperature. Grids were washed 3x 5 min each in Wash buffer, once in 0.1 M phosphate buffer, pH 7.4 and then mounted in Fluoromount anti-fade mounting compound. Bright field and fluorescent images were collected with a 10x, or 40x magnification using a 0.9 NA water immersion lens on a Zeiss inverted microscope using an Andor Zyla 4.0 CMOS camera. Exposure time, shutter and filter wheel (Sutter Instrument Co.) were controlled through Metamorph software (Molecular Devices).

### Cryo-electron Tomography

For tomographic data collection, single-axis tilt series were collected from -50° to +50° in 3° increments at approximately -8 □m under focus on an FEI Polara G2 operated at 300 kV equipped with a Gatan K2 Summit direct electron detector operated in photon counting mode. Data collection was performed in a semi-automated fashion using Serial EM software operated in low-dose mode [54]. Briefly, areas of interest were identified visually and 8 × 8 montages were collected at low magnification (2400x) and then individual points were marked for automated data collection. Data were collected at either 8.5 or 4.5 Å/pixel. Movies of 8-10 dose-fractionated frames were collected at each tilt angle and the electron dose spread across all images was limited to a total dose of < 100 e^−^/Å^2^ per tilt series. There is a “missing wedge” of information in the reconstructions due to the inability to collect tilt series through a full 180° (+/-90°) of stage tilting. Additionally, as the stage is tilted the electron path through the sample increases, decreasing the contrast and quality of the high tilt images. For our system, tilting the stage +/− 50° was found to be an optimal compromise. The missing wedge leads to anisotropic resolution producing elongation and blurring in the Z-dimension and tomographic reconstructions need interpreted acknowledging this limitation.

### Tomographic Reconstruction and Segmentation

Each tomographic data set was drift corrected with MotionCorr2 [55] and stacks were rebuilt and then aligned using IMOD [56, 57]. Tomograms of the aligned stacks were then reconstructed using TOMO3D [58, 59]. Contrast was enhanced using SIRT reconstruction implemented in TOMO3D.

Reconstructed tomograms were further processed using the median, non-local means, and Lanczos filters in Amira (FEI, ThermoFisher Scientific) for manual and semi-automated segmentation. Segmentation was accomplished by manually tracing membranes for each Z slice of the tomographic data set. Membranes were identified and segmented with reference to visualization in all three dimensions (X, Y, and Z). The brush tool was primarily used in manual segmentation. When possible, masking approaches were also used in combination with density thresholding for semi-automated segmentation. After segmentation, smoothing tools were employed for the manual tracings and surfaces were rendered for model construction. All measurements (length and diameter) were performed in either Amira using the 3D measurement tool or in IMOD.

## Acknowledgements

The authors would like to acknowledge Dr. Andrey Tsvetkov, Ndidi Uzor and Felix Moruno Manchon for help with primary neuronal cultures. We would also like to thank Dr. Michael Beierlein, Dr. Richard Youle and Dr. Matt Swulius for advice on the manuscript. This study was supported by grants from the National Institute of Health/National Institute of Neurological Disease and Stroke; R01NS101686 (MWN and PKD), R01NS087149 (PKD) and F31NS098790, Ruth L. Kirchstein National Research Service Award Predoctoral Fellowship (TF). PKD acknowledges support from the Nina and Michael Zilkha endowment and MNW acknowledges support from the William Wheless III professorship. The Polara electron microscope was supported, in part, through the Structural Biology Imaging Center at UTHSC—Houston. The Gatan K2 Summit was funded by the National Institutes of Health—Award S10OD016279.

## Supporting Figure Legends

Figure S1. 2D images from tomographic reconstructions of different membrane bound organelles observed in the varicosities of cryopreserved hippocampal neurons. A) mitochondrion, B) MVB and vesicles (arrowheads). C) ER and an unidentified membrane-bound compartment (arrowhead), D) Autophagosome, E) Lamellar body. Scale bar = 200 nm.

Figure S2. Different populations of mitochondrial cristae ultrastructures in varicosities. A) Representations of mitochondria displaying thin, tubulated cristae. B) Representations of mitochondria displaying thick, unstructured cristae. Populations were identified based on full 3D tomographic reconstructions. Scale bar = 200 nm.

Movie S1. Animation of the 3D tomographic reconstruction and overlay of the segmented structures in Figure 2. Scale bar = 200 nm.

Movie S2. Animation demonstrating space within an axon encountered by a mitochondrion presented in the X dimension. 3D tomographic reconstruction corresponds to Figure 3B and 5B. Scale bar = 200 nm.

Movie S3. Animation of the 3D tomographic reconstruction and overlay of the segmented structures in Figure 3A. 3D tomographic reconstruction also corresponds to Figure 5A. Scale bar = 200 nm.

Movie S4. Movie of the tilt series of tomographic data set corresponding to Figure 3A and 5A. Scale bar = 200 nm.

Movie S5. Movie of the 3D tomographic reconstruction corresponding to Figure 3B and 5B. Scale bar = 200 nm.

Movie S6. Movie of the tilt series of tomographic data set corresponding to Figure 3C and 5C. Scale bar = 200 nm.

Movie S7. Movie of a mitochondrion at spatially restricted axonal/varicosity boundary. Scale bar = 200 nm.

Movie S8. Movie of an additional mitochondrion at spatially restricted axonal/varicosity boundary. Scale bar = 200 nm.

Movie S9. Movie of the 3D tomographic reconstruction corresponding to Figure 4. Scale bar = 200 nm.

Movie S10. Movie of a 3D tomographic reconstruction displaying a mitochondrion spanning a short axon segment and residing in two varicosities. Scale bar = 200 nm.

Movie S11. Animation of the 3D tomographic reconstruction and overlay of the segmented membranes of mitochondria in Figure 3D. Scale bar = 200 nm.

